# A computational model of inhibition of HIV-1 by interferon-alpha

**DOI:** 10.1101/031005

**Authors:** Edward P Browne, Benjamin Letham, Cynthia Rudin

## Abstract

Type 1 interferons such as interferon-alpha (IFN*α*) inhibit replication of Human immunodeficiency virus (HIV-1) by upregulating the expression of genes that interfere with specific steps in the viral life cycle. This pathway thus represents a potential target for immune-based therapies that can alter the dynamics of host-virus interactions to benefit the host. To obtain a deeper mechanistic understanding of how IFNα impacts spreading HIV-1 infection, we modeled the interaction of HIV-1 with CD4 T cells and IFN*α* as a dynamical system. This model was then tested using experimental data from a cell culture model of spreading HIV-1 infection. We found that a model in which IFN*α* induces reversible cellular states that block both early and late stages of HIV-1 infection, combined with a saturating rate of conversion to these states, was able to successfully fit the experimental dataset. Sensitivity analysis showed that the potency of inhibition by IFN*α* was particularly dependent on specific network parameters and rate constants. This model will be useful for designing new therapies targeting the IFN*α* network in HIV-1-infected individuals, as well as potentially serving as a template for understanding the interaction of IFN*α* with other viruses.

**Author Summary:** Interferon-alpha (IFN*α*) is a key component of the host response to HIV-1, but the details of how IFN*α* regulates infection are still incompletely understood. To provide a deeper understanding of the dynamics of how IFN*α* inhibits HIV-1, we simulated the interaction of IFN*α* and HIV-1 as a computational model and compared this model to an experimental dataset. We identify a model structure that is able to fit many key features of the data. Furthermore, we use the model to predict optimal strategies for targeting the IFN*α* pathway therapeutically. We anticipate that this model will be useful for further analysis of HIV-IFNα interactions and will help to guide new therapeutic strategies.

## Introduction

Around 65 million people worldwide have been infected with Human immunodeficiency virus (HIV-1) [1]. Although progress has been made in mitigating disease with antiviral chemotherapy, a protective vaccine has proved elusive, and other approaches are still needed. Furthermore, permanently eliminating virus from patients undergoing drug therapy has been difficult due to the existence of latently infected reservoirs that are resistant to standard antiviral therapy [2]. Another possible approach to treating HIV-1 infection is to alter aspects of virus-host dynamics by targeting host pathways that inhibit or enhance infection. For this to be successful, a deep understanding of the dynamics underlying how specific host pathways interact with HIV-1 will likely be required.

The application of mathematical modeling to HIV-1 dynamics during acute and chronic infection has been highly successful at improving our understanding of the basic features of clinical infection. In particular, fundamental insights into the response to antiviral therapy, and the existence of multiple long-lived virus reservoirs have been revealed [3–6]. In early models, the extent of infection was typically limited by target cell abundance, although more recent HIV-1 models have also considered the impact of virus-specific CD8 T cells [7,8]. However these models have not yet included the impact of the host innate immune system.

A key component of the host innate response to HIV-1 infection is the type 1 interferon (IFN) system [9,10]. In humans, type 1 IFNs consist of a family of related cytokines including 13 subtypes of IFNα, and two subtypes of IFNβ, that are secreted in response to stimulation of microbe-sensing pattern-recognition receptors such as Toll-like receptors (TLRs), RIG-I-like receptors (RLRs) and NOD-like receptors (NLRs) [11]. Type 1 IFNs bind the IFN*α* receptor (IFNΑR) and actevate phosphorylation of the signaling molecules STAT1 and STAT2, which then bind to Interferon regulatory factor 9 (IRF9) to form the Interferon-stimulated gene factor 3 (ISGF3) complex [12]. ISGF3 then binds to conserved Interferon-sensitive response elements (ISREs) found upstream of interferon-sensitive genes (ISGs) [13]. Dozens of ISGs are upregulated by IFN*α*, the function of which only a few are clearly understood [14]. Overall, IFNs create an antiviral state that can either prevent *de novo* infections, or inhibit later stages of virus replication in cells, such as assembly and egress.

IFNα is detectable in the plasma during acute HIV-1 infection, and this cytokine is predominantly secreted by plasmacytoid dendritic cells (pDCs). pDCs detect HIV-1 via the single-stranded RNA sensor TLR7, and secrete high levels of IFNα due to constitutive expression of the Interferon regulatory factor 7 (IRF7) transcription factor [15,16]. A number of ISGs have been shown to have anti-HIV-1 activity [17]. In particular, Tripartite motif-containing 22 (TRIM22) and Myxovirus resistance protein 2 (MX2) inhibit early stages of infection [18,19], while other ISGs such as Apolipoprotein B mRNA editing enzyme, catalytic polypeptide-like 3G (APOBEC3G) and Tetherin target later stages of infection, such as virus release or the infectivity of virus particles [20,21]. Furthermore, IFNα inhibits HIV-1 replication in tissue culture and blockade of IFNα during Simian immunodeficiency virus (SIV) infection leads to higher virus levels *in vivo* [10].

Due to its HIV-1-inhibiting properties, IFNα has attracted interest as a therapeutic target for HIV-1 infection. However, treatment of HIV-1 patients with recombinant IFNα has produced inconsistent and disappointing results, with only modest effects on virus levels being observed [22–24]. However, since the structure of the IFNα inhibitory network, as well as the parameters that regulate its activity are poorly understood, a better understanding of the underlying dynamics of this response may lead to improved and more effective IFNα-based therapies. For example, dynamical models may help pinpoint which molecular components of the IFNα inhibition network will achieve the most potent or most durable results. Furthermore, modeling approaches have previously been applied to the interaction of Hepatitis C virus (HCV) with peggylated IFNα, and have yielded valuable insights [25–27].

To achieve this goal, we simulated the inhibition of HIV-1 by IFNα using a dynamical system modeling approach, and tested this model in a well-defined experimental system. In our model, IFNα interacts with both HIV-1-infected and uninfected cells to induce a reversible state of blocked infection, and we demonstrate that this model makes testable predictions about how specific network parameters may be targeted to achieve maximal inhibition of HIV-1.

## Results

### IFNα inhibits spreading HIV-1 infection

To generate an experimental dataset against which different models can be tested, we performed experiments with a tissue culture model of spreading HIV-1 infection. CEM-5.25 cells are a human CD4 T cell line that are susceptible to HIV-1 infection and contain an integrated HIV-1 long terminal repeat-Green fluorescent protein (LTR-GFP) reporter cassette [28]. Infected cells express GFP due to the viral Tat protein permitting transcription of GFP from the integrated LTR. This allows us to distinguish HIV-1-infected cells (GFP+) from uninfected cells (GFP−). CEM-5.25 cells were infected with HIV-1, and both cell and supernatant samples were then isolated at 24h intervals for 72h post infection. Cells samples were analyzed by flow cytometry to determine 1) absolute cell numbers and 2) the percentage of HIV-1-infected (% GFP+) cells. Infectious virus in the supernatant was quantified by focus-forming assay using GHOST-X4 reporter cells [29]. In infected cell cultures, the *%* GFP+ cells, as well as the concentration of infectious HIV-1 in the supernatant, rises exponentially until 3dpi (Fig. 1B,C). By day 3, total cell levels decline due to infection-induced cytopathic effect (Fig. 1A).

**Figure 1:**
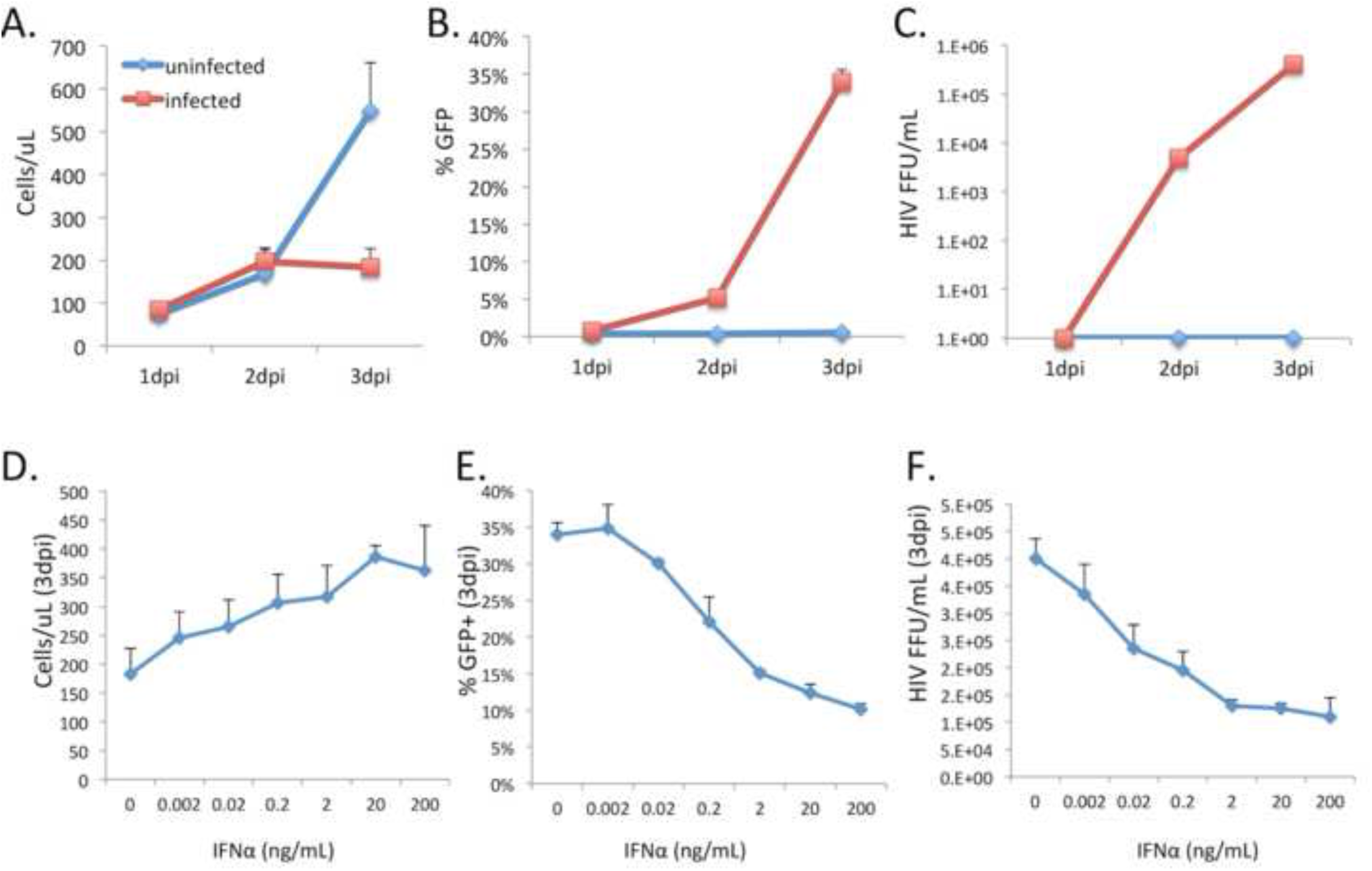
Inhibition of spreading HIV-1 infection by IFNα. A human CD4 T cell line was infected with HIV-1 and the total concentration of cells (A), the proportion of infected (GFP+) cells (B), and the concentration of infectious virus in the supernatant (C), were monitored at 24h intervals. Different concentrations of IFNα were added to the cells at 6h prior to infection, and the effect on total cell concentration (D), the percent infected cells (E), and the concentration of infectious HIV-1 in the supernatant (F), were measured every 24h. Measurements were taken in quadruplicate and data shown are representative of three independent experiments. Error bars represent the standard deviation of the dataset.

To examine the effect of IFNα on the replication of HIV-1 in this system, different concentrations of IFNα were included in the tissue culture media from 6h prior to infection and maintained throughout the course of the infection. Inclusion of IFNα in the media caused a clear and progressive decrease in the accumulation of infected cells (Fig. 1E) and infectious HIV-1 in the supernatant (Fig. 1F), as well and an increase in overall cell density at later timepoints (Fig 1D). Interestingly, inhibition of HIV-1 by IFNα exhibits two key features – firstly that the inhibition curve is very broad – with differential inhibition being observed for IFNα concentrations over several orders or magnitude (from 2pg/mL to 20ng/mL), and secondly, that inhibition of HIV-1 increased only minimally above 2ng/mL, suggesting that inhibition saturates at higher IFNα concentrations. 50% inhibition of infection, as measured by infectious HIV-1 concentration at 3dpi, was achieved between at 0.02ng/mL and 0.2ng/mL (Fig. 1F).

### Model of IFNα inhibitory network

We constructed a dynamical system that models the dynamics of CD4 T cells and their interaction with both HIV-1 and IFNα (Fig. 2, Table 1). This model shares a number of features with a model previously used to analyze the IFNα response to influenza infection [30]. In this system, the initial species are HIV-1 (H), CD4 T cells (C) and IFNα (I). HIV-1 can infect CD4 T cells to generate infected cells (CH) at a rate proportional to the concentrations of HIV-1 and CD4 T cells, *via* the infection rate constant (k_5_). Infected cells secrete infectious HIV-1 (H) at a constant rate (k_6_). IFNα can bind to uninfected (C) or infected cells (CH) and convert them into refractory cells (CI) that cannot be infected, or infected-inhibited cells (CHI) that no longer release virus, respectively. The rate of this conversion is determined by the concentrations of susceptible cells and IFNα, as well as the k_2_ and k_8_ rate constants. Since the effects of IFNα on cells are reversible, CI and CHI revert at an intrinsic rate (k_3_) to C or CH respectively. H and I each have their own intrinsic decay rates that were measured separately (k_7_ and k_9_) (Fig. S1). For these studies, an immortalized CD4 T cell line (CEM 5.25) was used that divides continuously and has an intrinsic death rate that is negligible when maintained at subconfluency. By contrast, HIV-1-infected cells (CH and CHI) cells die at an intrinsic rate due to the cytopathic effect of HIV-1 infection (k_4_). All CD4 T cell subsets divide at an intrinsic rate to generate additional cells of the same type. The replenishment rate of all CD4 T cells was modeled with the same rate for all subspecies (k_1_), which was experimentally determined by parameter fitting.

**Figure 2:**
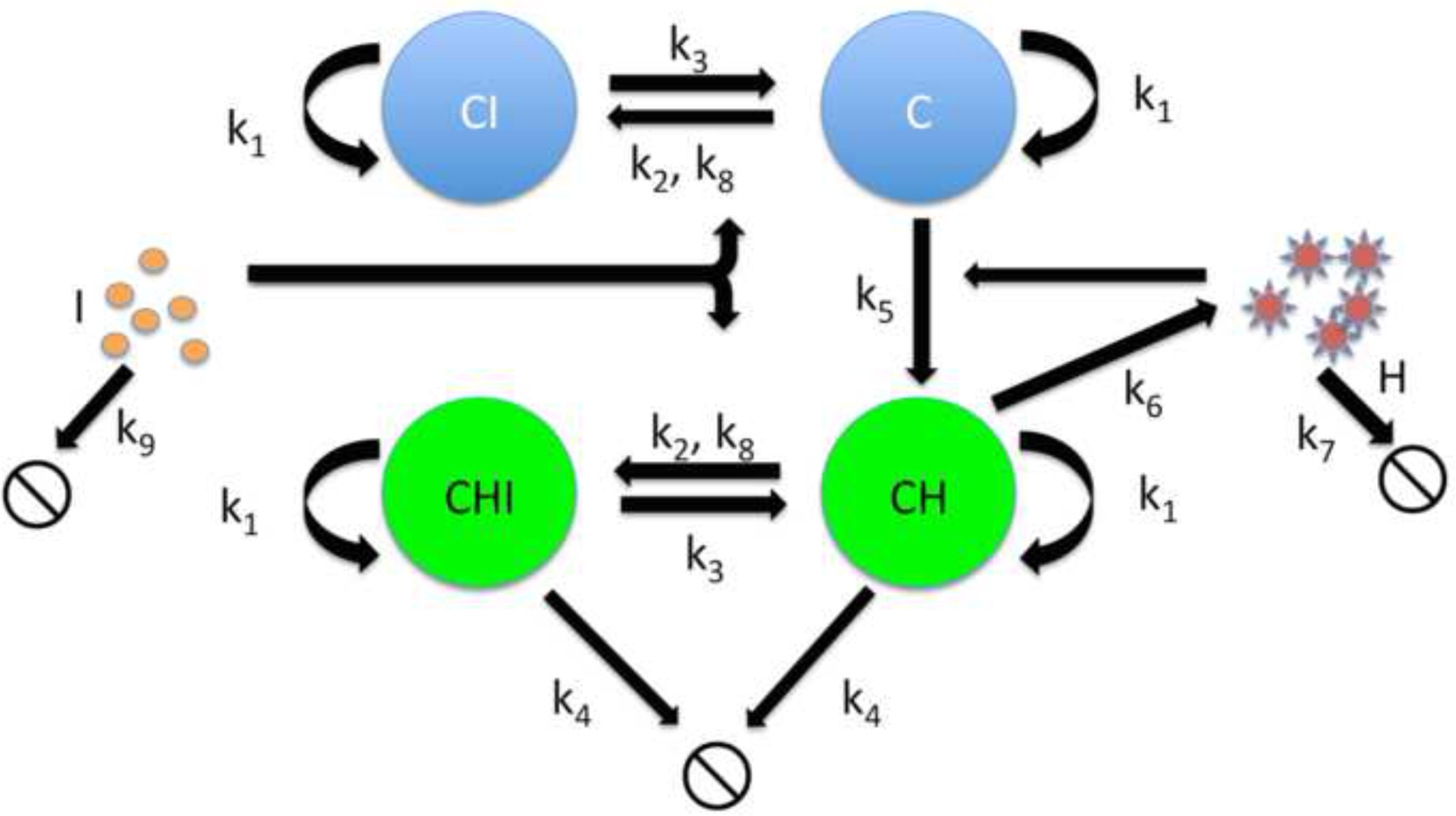
Model of inhibition of HIV-1 by IFNα. CD4 T cells are replenished at a rate proportional to the number of cells, leading to exponential growth (k_1_). HIV-1 particles infect cells (k_5_), and infected cells die at an inherent rate (k_4_). IFNα inhibits infection by converting uninfected cells (C) and infected cells (CH) into states in which they cannot be infected (CI), or which longer release infectious virus (CHI) (k_2_), and this rate is also governed by a saturating rate constant (k_8_). Both these states are reversible, and without continued IFNα exposure, CI and CHI revert to C and CH respectively (k_3_). Infected cells (CH) secrete HIV-1 at a continuous rate (k_6_). Both HIV-1 and IFNα have their own natural decay rates (k_7_ and k_9_ respectively).

**Table 1:** Species and parameter descriptions.

| Species/parameter | description | Units |
| --- | --- | --- |
| C | Uninfected CD4 T cells | cells $\mu\text{L}^{-1}$ |
| CI | Uninfected CD4 T cells made refractory to infection by IFNα | cells $\mu\text{L}^{-1}$ |
| CH | HIV-1-infected CD4 T cells | cells $\mu\text{L}^{-1}$ |
| CHI | HIV-1-infected CD4 T cells in which virus release | cells $\mu\text{L}^{-1}$ |
| | has been blocked by IFNα | |
| H | HIV-1 | Focus-forming units (FFU)<br>uL<sup>-1</sup> |
| I | Interferon alpha | ng mL<sup>-1</sup> |
| k<sub>1</sub> | Intrinsic division rate of CD4 T cells | day<sup>-1</sup> |
| k<sub>2</sub> | Rate of conversion of CD4 T cells to refractory (CI)<br>or blocked (CHI) state. | ng<sup>-1</sup> day<sup>-1</sup> |
| k<sub>3</sub> | Intrinsic rate of reversion of IFNα exposed cells to<br>HIV-1 susceptible state. | day<sup>-1</sup> |
| k<sub>4</sub> | Death rate for HIV-1 infected cells (CH and CHI) | day<sup>-1</sup> |
| k<sub>5</sub> | Infection rate | uL day<sup>-1</sup> FFU<sup>-1</sup> |
| k<sub>6</sub> | HIV-1 secretion rate from infected cells (CH) | FFU cell<sup>-1</sup> day<sup>-1</sup> |
| k<sub>7</sub> | HIV-1 decay rate | day<sup>-1</sup> |
| k<sub>8</sub> | Saturated rate of conversion to CI or CHI | ng mL<sup>-1</sup> |
| k<sub>9</sub> | IFNα activity decay rate | day<sup>-1</sup> |

One key element of our model was the inclusion of a Michaelis-Menten saturating rate constant (k_8_) for the conversion of C to CI and CH to CHI. As such, the conversion rate of C to CI and CH to CHI saturates for high I concentrations, due to k_8_. Previous models of IFNα interaction with virus infection do not include saturation for conversion to an inhibited state [30], and this therefore represents a novel feature of this model.

Our system of equations is thus:

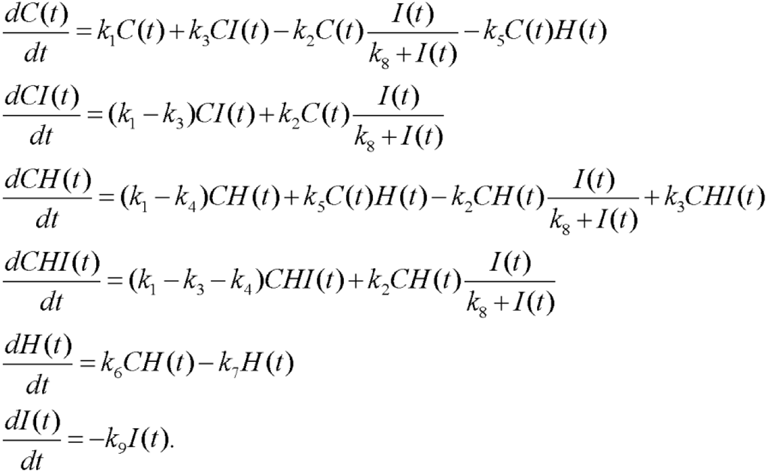

### Derivation of parameter estimates

First, the biological decay rates for HIV-1 (k_7_) and IFNα (k_9_) were experimentally determined separately and assumed to be invariant in the different conditions (Fig. S1). We then estimated all seven remaining parameters (k_1_ through k_6_ and k_8_) and the initial CD4 T cell count C_(0)_ using the 72h timecourse data. In particular, we estimated these parameters by minimizing the weighted sum of squared distances between each observation and the corresponding model prediction. Using our experimental data, we cannot directly distinguish between individual uninfected subspecies (C and CI) or infected subspecies (CH and CHI). Thus, the observed cell species, GFP+ and GFP− cells, were assumed to represent the sum of CH and CHI, and the sum of C and CI, respectively.

We compared the basic model (Model 1) to several variants of the model to determine which could more accurately fit the dataset, and to identify the features that make important contributions to model accuracy. For Models 1, 3 and 4, the rate of conversion to inhibited cell states saturates due to the inclusion of the k_8_ rate constant, while for models 2, 5 and 6 we take the conversion rate of C to CI and CH to CHI as linearly proportional to I, as has been done in previous models of IFNα interaction. For Model 3, cell division was restricted to uninfected CD4 T cells that had not been exposed to IFNα (C). For Model 4, we only considered the interaction of IFNα with uninfected CD4 T cells, and ignored inhibition of previously infected cells – meaning that the CHI species was eliminated from the model. For Model 5, the death rate (k_4_) for the CHI species was modeled as distinct from the death rate of CH, while for Model 6, the interconversion rate constants (k_2_ and k_3_) between uninhibited and inhibited states were distinct for infected and uninfected cells. Different model configurations are described graphically in Figure S2.

The accuracy of the model fits were compared between these six different model configurations (Table 2). Notably, we found that inclusion of the saturation constant for IFNα-mediated inhibition dramatically improved the accuracy of the fitting, with significantly lower fit error for all model variants with saturating inhibition (Models 1, 3 and 4). Model 1 most accurately simulated overall timecourse data, and became our default model for further analysis. This model was also significantly superior to another variant with saturating inhibition that lacks CHI (Model 4) suggesting that considering the effects of IFNα on infected cells is important. Model 1 also accurately simulated important features of the timecourse data for %GFP− and GFP+ T cells, as well as the concentration of infectious HIV-1 in the supernatant (Figure 3A-C). The better fit for the saturating models was not simply due to the inclusion of an extra rate constant (k_8_), because other variants of the linear inhibition model with additional parameters (models 5 and 6) did not exhibit a comparable improvement in fit quality.

**Table 2:** Sum of squared error for model fit versus measured species. A nonlinear least squares method was used to fit experimental data to six different model configurations – some with saturating inhibition by IFNα (Models 1, 3 and 4), and others with linear inhibition (Models 2, 5 and 6). Weighted squared error (Fit error) was determined for the best fit for each model with respect to the all measured concentrations of GFP− cells, GFP+ cells, and HIV-1. Fit error total refers to the overall mean error of the fit to all collected data. For each model, a 95% confidence interval for the increase in error over that of Model 1 (95% CI), and the Akaike Information Criterion (AIC), are shown. An asterisk next to the fit error value for a model indicates a statistically significant difference in performance relative to Model 1, at the *p*<0.05 level.

| Model | Description | Parameters | Fit error | 95% CI | AIC |
| --- | --- | --- | --- | --- | --- |
| 1 | Basic model (saturating inhibition). | 8 | 277.5 |  | 2529 |
| 2 | Linear inhibition | 7 | 849.5* | 240.7 - 984.0 | 3671 |
| 3 | Saturating inhibition, cell division restricted to C species | 8 | 296.6 | -1.7 - 42.3 | 2567 |
| 4 | Saturating inhibition, no effect of IFNα on infected cells (No CHI) | 8 | 297.7* | 6.6 - 35.5 | 2570 |
| 5 | Linear inhibition, separate death rates ($k_4$) for CH and CHI | 8 | 848.4* | 240.1 - 985.3 | 3671 |
| 6 | Linear inhibition, separate interconversion rates ($k_2$ and $k_3$) for uninfected (C and CI) and infected (CH and CHI) cells. | 9 | 846.0* | 240.1 - 976.3 | 3669 |

**Figure 3:**
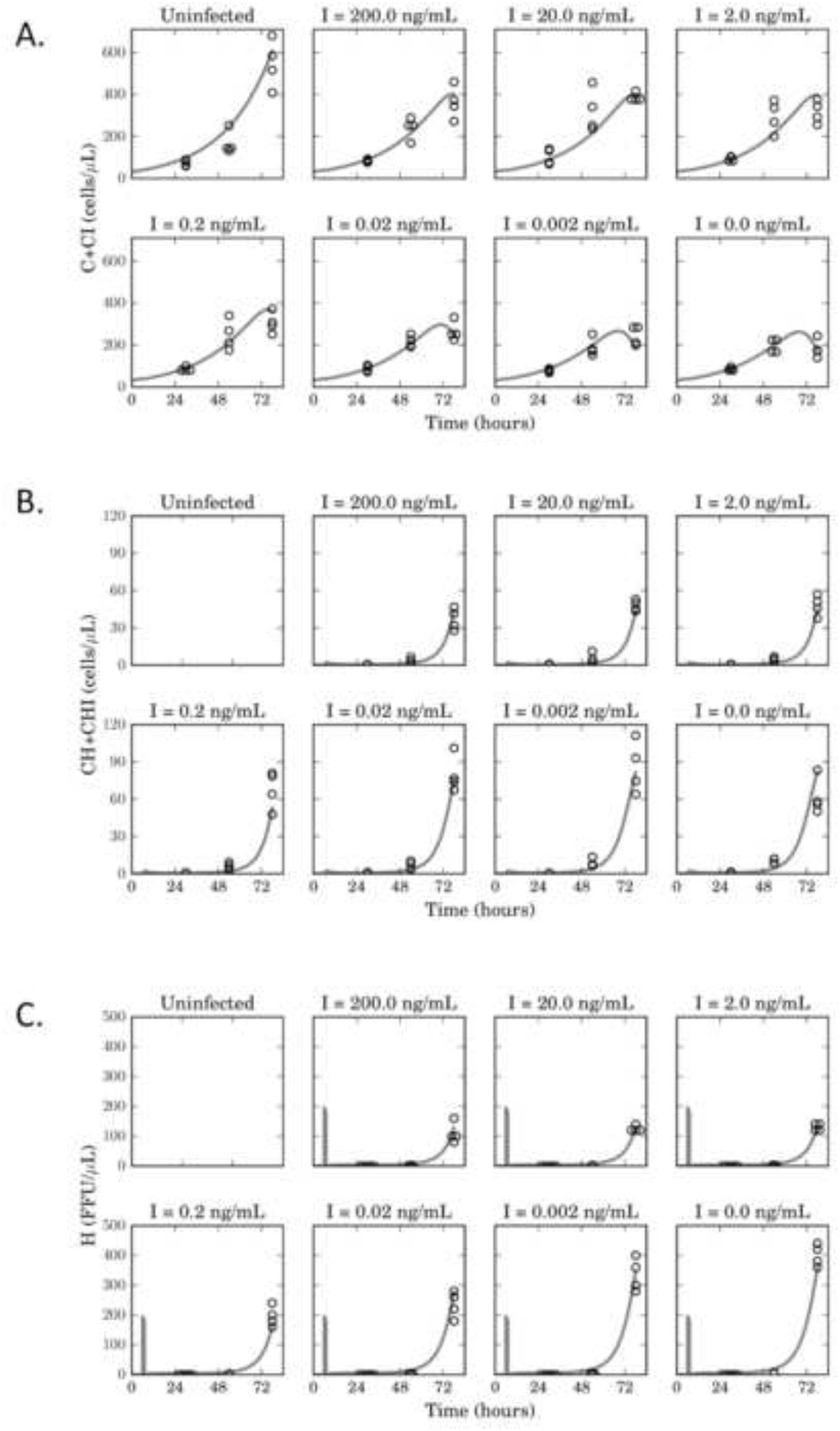
A saturating model of inhibition of HIV-1 by IFNα recapitulates infection timecourse data. Experimental data for the concentration of uninfected (GFP-) cells (A), infected (GFP+) cells (B), and infectious HIV-1 (C) was fitted with a saturating model (Model 1). Open circles represent independent experimental data points, while for each species, the solid black lines represent the model’s best fit. Data points with the same value are shown side by side.

Estimated parameter values for the model rate constants were determined for all six model variants (Table 3, Table S1). Estimates for most parameters except k_2_, k_3_ and k_8_ were similar between Model 1 (saturating) and Model 2 (linear). However, compared to Model 1, Model 2 exhibited a significantly lower k_2_ estimate and a higher k_3_ estimate. Also, the predicted concentrations for each individual CD4 T cell species were calculated from the best fit for both Model 1 and Model 2. Notably, for most concentrations of IFNα, Model 1 predicts a higher proportion of cells in the inhibited states compared to Model 2, although both models predict that the majority of inhibited cells are uninfected over the range of IFNα concentrations tested (Fig S4).

**Table 3:** Estimated model parameter values. Values for unfixed rate constants were obtained as the best fits of Model 1 and Model 2 to the experimental dataset.

| Parameter | Units | Model 1 value (95% CI) | Model 2 value (95% CI) |
| --- | --- | --- | --- |
| k1 | day<sup>-1</sup> | 0.905 (0.831-1.006) | 0.898 (0.831-0.945) |
| k2 | ng<sup>-1</sup> day<sup>-1</sup> | 37.5 (31.5-45.1) | 0.0431 (0.0291-0.0610) |
| k3 | day<sup>-1</sup> | 261 (214-552) | 355 (0-inf) |
| k4 | day<sup>-1</sup> | 6.42 (4.80-8.61) | 8.13 (5.86-10.96) |
| k5 | uL day<sup>-1</sup> FFU<sup>-1</sup> | 0.00913 (0.00736-0.0116) | 0.0136 (0.0104-0.0175) |
| k6 | FFU cell<sup>-1</sup> day<sup>-1</sup> | 20.1 (18.3-23.1) | 14.4 (13.3-15.2) |
| k8 | ng mL<sup>-1</sup> | 0.154 (0.063-0.463) | NA |

To further investigate the performance of our model, we re-plotted the data for HIV-1 levels, % GFP+ cells, and total cell concentration, at 3dpi, against different IFNα concentrations, and compared to model predictions (Fig. 4). Significantly, Model 1 was able to simulate key features of the HIV-1 inhibition curve accurately including plateaus at high and low IFNα levels, and a broadly sloped inhibitory curve. Furthermore, Model 1 performed better than a linear inhibition model (Model 2) in two out of the three curves, with statistically significantly lower RMSE values for HIV-1 concentrations and the percentage of infected cells.

**Figure 4:**
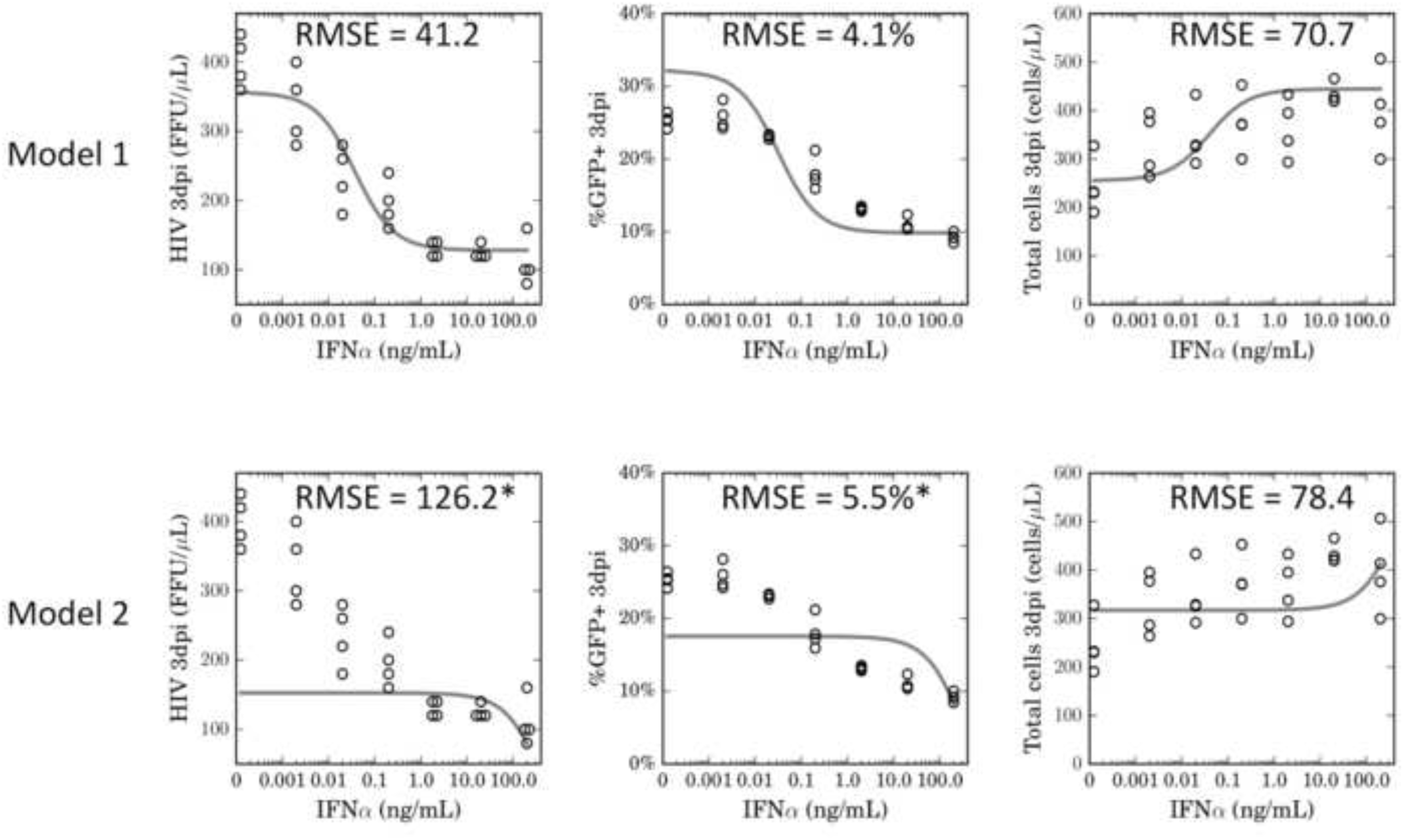
Saturating effect of IFNα on CD4 T cells improves model accuracy. Dose-response data for the effect of different IFNα concentrations on the concentration of HIV-1 (left panel), the percentage of GFP+ cells (middle panel), and on total cell concentration (right panel) from the 3dpi timepoint of the timecourse dataset were compared to model generated estimates for Models 1 and 2. Root mean squared errors (RMSE) are shown on the figure panels. Circles represent individual data points. An asterisk besides RMSE values for Model 2 denotes a statistically significant difference in performance relative to Model 1.

### Differential effects of inhibitory parameters on the potency of IFNα

Using the saturating default model, we examined the sensitivity of biologically significant outcomes of the model to specific parameters - particularly the parameters that regulated the IFNα-related component of the network (k_2_, k_3_ and k_8_). To measure the outcome of parameter scanning, we calculated the HIV-1 concentration at 3dpi as a function of IFNα concentration with different parameter values over a range of 10 fold higher or lower than our estimated values. As expected, increasing the rate at which IFNα converts CD4 T cells to an inhibited state (k_2_), and decreasing the reversion rate (k_3_), both individually resulted in lower total HIV-1 amounts at most concentrations of IFNα (Fig 5A). Modulating the saturation rate constant (k_8_) had negligible effect on HIV-1 levels at higher IFNα concentrations, but modulated the IFNα threshold at which saturation is achieved. Thus, although the inclusion of a saturation constant greatly improved accuracy of the model, the outcome of infection is less sensitive to the value of this parameter than for k_2_ and k_3_ at higher IFNα concentrations.

To describe the effects of IFNα on HIV-1, two metrics have previously been used – the IC_50_, that is, the IFNα concentration that results in 50% of maximal inhibition, and the V_res_, which is the percent residual HIV-1 replication at the maximal IFNα concentration [31]. Based on our simulations, we observe a clear differential effect of k_2_, k_3_ and k_8_ on and V_res_, namely that k_2_ and k_3_ strongly affect V_res_, while k_8_ does not (Fig 5B). Interestingly, increased k_2_ values and decreased k_3_ values both reduced the IC_50_ of IFNα, while scaling these parameters in the opposite direction had little effect. By contrast, both higher and lower k_8_ values affected the IC_50_, with lower k_8_ leading to a lower IC_50_, and higher k_8_ leading to a higher IC_50_. Overall, we conclude that V_res_ is determined by the balance of k_2_ and k_3_, while IC_50_ is determined by the combined effect of k_2_, k_3_ and k_8_. Thus our model makes testable predictions about how these specific network parameters affect IFNα inhibition of HIV-1.

**Figure 5.**
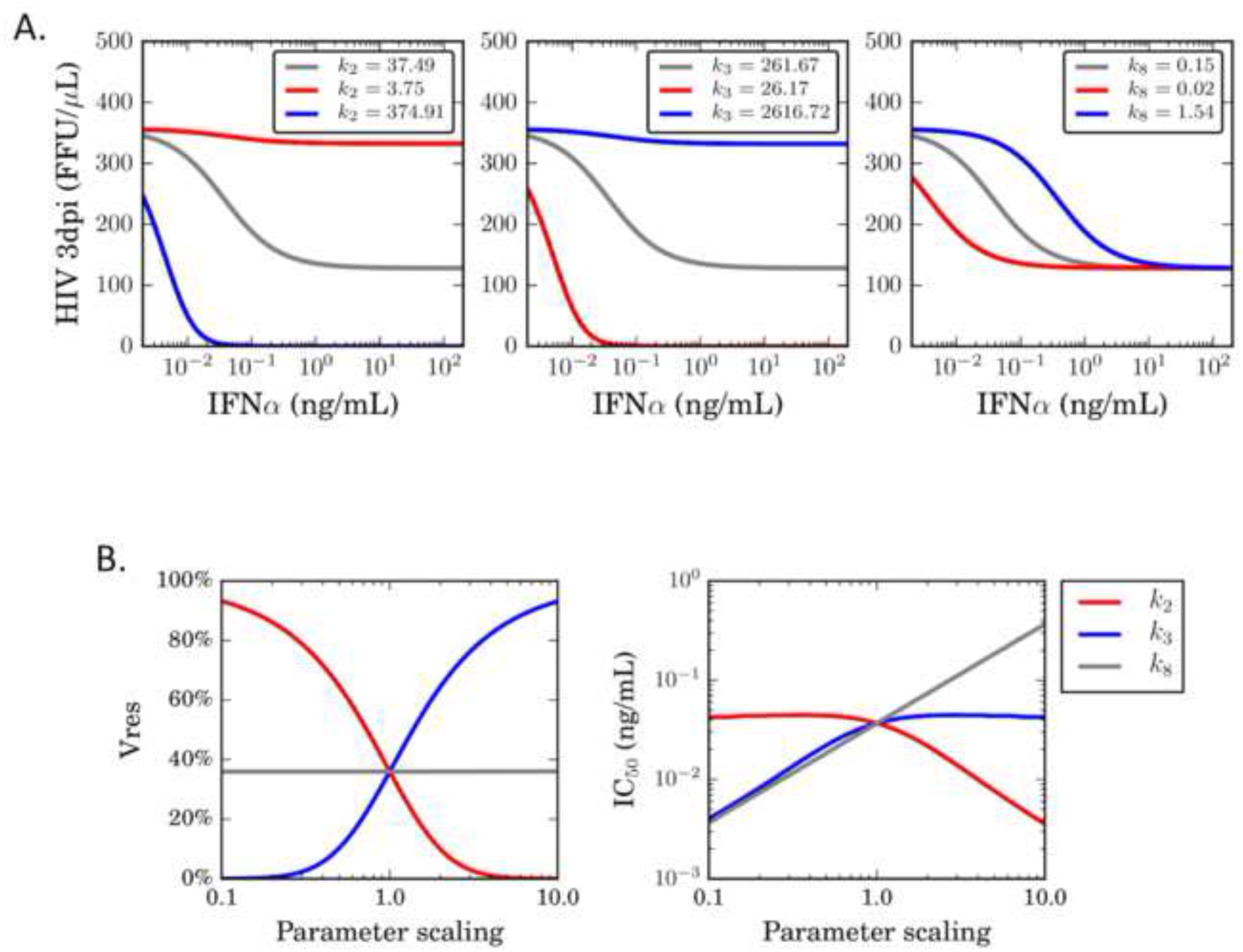
Differential effects of inhibitory parameters of the potency of IFNα. (A) Experimentally measured values for k_2_ (left panel), k_3_ (middle panel) and k_8_ (right panel), were modulated up (blue) and down (red) by 10-fold, and the effect on the outcome of infection determined by running a simulation with these parameter values substituted for the experimentally determined ones, and calculating the concentration of infectious HIV-1 at 3dpi. (B) The effect of modulating values for k_2_, k_3_ and k_8_ on IC_50_ (concentration required for 50% inhibition) and V_res_ (the percent HIV-1 replication at maximal IFNα concentration) were determined by scanning parameter values over a 10 fold range above and below our measured value, and calculating the HIV-1 concentration at 3dpi.

## Discussion

In this manuscript we describe, for the first time, a dynamical model that simulates the interaction of HIV-1 with IFNα, and demonstrate that this model accurately fits an experimental dataset. Also, we have estimated rate constants for how IFNα interacts with CD4 T cells and HIV-1, thereby providing a realistic range in which to perform simulations of infection. A key novel feature of this model is a saturating rate constant for IFNα’s effects on target cells, which significantly increased the accuracy of the model’s fit to the experimental data. In the absence of this feature, the model performed poorly in fitting, and had a profoundly different balance of forward and reverse rate constants for the generation of inhibited cellular states (k_2_ and k_3_). Furthermore, we have found that a model in which IFNα affects both uninfected and infected cells fit experimental data significantly better that one in which IFNα only affected uninfected cells. Although the interaction of IFNα with viral infection has been previously studied using modeling approaches [30,32–34], a saturating rate constant for inhibition of infection has not been a standard assumption of these models. A similar study that considered inhibition of influenza by IFNα found that a linear inhibition model, in which IFNα affects uninfected cells only, was able to successfully fit experimental data, suggesting that models of IFNα interaction with viruses may require virus-specific dynamics [30].

This model could be useful for the design of novel therapies for HIV-1 that target the IFNα pathway. Notably, our model recapitulates the observation that IFNα has a broad HIV-1-inhibition curve, meaning that changes in IFNα concentration over several orders of magnitude have only partial effects on HIV-1 levels. However, sensitivity analysis of the network rate constants indicates that modulation of the k_2_ and k_3_ rate constants can lead to a dramatically more potent inhibition of HIV-1 for a given IFNα concentration. By contrast, the value of the saturation rate constant (k_8_) has a lesser effect on the course of infection for most IFNα concentrations. As such, our model predicts that clinical modulation of k_2_ and k_3_ could prove more beneficial than simply boosting total IFNα levels by a similar factor or by modulating k_8_.

In order to design therapies that can target the biological processes controlled by these rate constants, it will be critical to identify the molecules that govern them. The signaling pathway and transcriptional response to IFNα have been extensively characterized, and the identities of many of the participating molecules have been described [14]. The rate of conversion to inhibited states (k_2_) could reflect features such as the abundance of the IFNAR receptor, the expression level of IFNα signaling factors such as STAT1/2 and IRF9, as well as the transcription/translation rate for antiviral effector proteins. The reversion rate (k_3_), by contrast, is likely determined by the off-rate for ISGF3 detachment from the IRSE sequences in the promoters of ISGs, and/or the degradation rate for antiviral effector proteins in the cytoplasm. The requirement for a saturating rate constant (k_8_) to accurately fit our dataset could reflect any rate-limiting process in the generation of an ‘inhibited’ cell. Factors that contribute to these rate constants could potentially be experimentally identified by manipulating expression of known components of this pathway, and examining the impact on the network behavior. Novel therapies designed to target these rate constants could involve boosting or reducing the expression/activity of these key factors.

The application of this model to clinical data from HIV-1 patients could yield valuable insights into HIV-1 immunity and pathogenesis. HIV-1 infection results in a wide range of outcomes in terms of viral loads, immune responses, and disease progression. If parameter values can be derived for individual patients, these parameters could be examined for correlation with any of these clinical outcomes. For example, do non-progressors and rapid progressors have different parameter sets that can potentially contribute differences in the course of clinical infection? Some evidence suggests that this may indeed be the case - females mount an intrinsically stronger IFNα response than males, and also exhibit stronger early control of HIV-1 [35–38]. The dynamics of early acute HIV-1 infection, and the resulting innate immune response, likely contains valuable information relating to the IFNα response of individual patients that could be analyzed by a version of this model, but unfortunately, data from this phase of infection have been difficult to obtain, since most patient are not diagnosed until they have entered the chronic phase of infection.

An important consideration for IFNα-related therapy is that IFNα may play different and potentially opposing roles at different times during HIV-1 infection. Recent data suggest that while IFNα and pDCs do indeed limit HIV-1/SIV levels during early infection, they may also promote CD4 T cell depletion at later times, possibly due to IFNα-induced upregulation of apoptosis [39]. However, administration of high doses of IFNα to SIV-infected African green monkeys, which naturally tolerate infection, does not enhance CD4 T cell loss or pathogenesis [40]. Furthermore, blockade of IFNα signaling in SIV-infected Rhesus Macaques leads to higher viral loads and more rapid disease progression [10]. Therefore, it is unclear if future models of IFNα’s role in clinical infection will have to incorporate this feature of IFNα’s behavior in order to accurately model the disease process.

Our model should also be considered in the light of some inherent caveats. For example, the model relies on mass-action interactions to simulate the innate immune response to infection. While this may accurately model responses in well-mixed compartments such as the blood, it may not be applicable to immune responses in solid tissues such as lymph nodes or mucosal surfaces. IFNα may also have additional indirect mechanisms of influencing the course of infection, such as helping to promote immune responses by Natural killer (NK) cells or lymphocytes [41,42]. Also, the results of this manuscript rely on an immortalized cell line that may behave differently than primary human CD4 T cells *in vivo.* Furthermore, HIV-1 employs several countermeasures against inhibitory ISGs, and different virus strains may exhibit differential sensitivity to IFNα [31]. Currently this model does not consider the contribution of endogenous IFNα to the course of infection. *In vivo,* the endogenous IFNα response is driven by plasmacytoid DCs (pDCs), and future development of this model to analyze clinical data from HIV-1 patients will likely require the incorporation of pDCs and endogenous IFNα. Our model also makes several assumptions that may not fully reflect details of HIV-1 infection, such as assuming that infected and uninfected CD4 T cells have equivalent division rates.

Nevertheless, this study represents the first attempt to analyze the interaction of the innate immune system with HIV-1 from a computational perspective, and demonstrates that quantitative estimates for the parameters that regulate IFNα’s potency can be derived from experimental data. As such, these findings should serve as a valuable starting point for future studies investigating the dynamics of the host innate immune response to HIV-1.

## Materials and methods

### Cells and viruses

CEM 5.25 cells and GHOST-X4 cells were obtained from Dan Littman (NYU). These cells were maintained in RPMI or DMEM media respectively with 10% fetal calf serum, glutamine, and penicillin/streptomycin. An integrated LTR-GFP cassette present in the genomic DNA of these cells facilitates expression of Green Fluorescent Protein (GFP) upon HIV-1 infection.

The HIV-1 strain used was NL4-3, a CXCR4-tropic subtype B strain [43]. The pNL4-3 plasmid was obtained from aidsreagent.org. To generate stocks of infectious virus, 10ug of plasmid was transfected into subconfluent 10cm plate of HEK-293FT cells (Invitrogen) using Mirus LT-1 reagent (Mirus Bio). At 4hrs post-transfection, medium was replaced. At 2 days post-transfection, the supernatant was harvested, clarified by low-speed centrifugation, then filtered through 0.45uM filter. The samples were then aliquoted and frozen at −80C. The titer of the stock was the determined by focus-forming assay.

For experimental infections, CEM-5.25 cultures were first exposed to IFNα for 6 hrs, then infected with NL4-3 at a multiplicity of 0.1 focus-forming unit per cell for 1hr. The virus inoculum was then removed and replaced with fresh media containing IFNα. The infected cells were then plated in 96 well plates at approximately 100 cells/uL. At 24h intervals a small fraction of the media (10%) was removed and replaced with media containing fresh IFNα.

### Focus forming assays

GHOST-X4 cells were plated at 10000 cells per well in a 96 well plate. A ten-fold dilution series of each supernatant sample was made and 100ul of each dilution was used to innocculate GHOST-X4 cells. At 3dpi, the supernatant was removed and the cells were fixed in 4% paraformaldehyde (PFA). The plate was then analyzed by fluorescence microscopy and the number of GFP+ foci per well was counted. This was then used to calculate the concentration of focus-forming units (FFU) per mL in each of the original supernatant samples.

### Flow cytometry

Cell samples from infected CEM-5.25 cultures were fixed in 2% PFA for 20 minutes, then diluted in phosphate buffered saline (PBS) and analyzed on an Accuri C6 flow cytometer. The total concentration of CD4 T cells, and the percentage of infected cells (% GFP+), was calculated for each original sample.

### Interferon

Human IFNα2a was obtained from Sigma Aldrich, reconstituted in PBS, and stored at - 80C.

### Computational analyses

Model fitting, simulation, and other computational analyses were done using the Python programming language. The system of differential equations was integrated using SloppyCell [43]. Model fitting was done by solving a weighted nonlinear least squares problem. Let *x_ijlt_* represent the data collected for species *i* at interferon level *j,* in replicate *l* at time point *t*. Let *y_ijt_(k)* be the corresponding model prediction for that species, interferon level, and time point, with parameters *k*. The weighted least squares problem is to minimize the fit error:

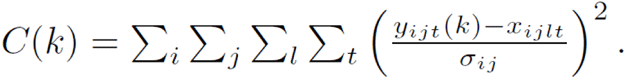

The measurement variance, 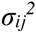, was estimated by first measuring the variance across replicates at each time point, and then averaging these variances over time points. The nonlinear squares problem was solved using random-restart optimization. For each random-restart run, optimization was doing using the L-BFGS-B optimization routine [44,45]. Optimization was started from 25 randomly selected initial values, and then the final parameters were chosen as those that produced the lowest cost across all 25 runs.

Confidence intervals for the best-fit parameters were found using the profile likelihood method [46]. The parameter confidence intervals reported here are simultaneous 95% confidence intervals.

Model comparison was done using two complementary methods: AIC and nonparametric confidence intervals on fit error. AIC provides a model comparison that incorporates the number of parameters, but requires assuming a normal noise model. Nonparametric confidence intervals were estimated using the percentile bootstrap, so do not require any assumption about the noise distribution. To determine if a difference in fit error between two models was statistically significant at the p<0.05 level, 95% confidence intervals were computed for the difference in fit errors. If this 95% confidence interval contained zero, it indicated that the difference in model fit was not statistically significant at the 0.05 level. If the confidence interval did not contain zero, it indicated that the difference in model fit was statistically significant.

---

**Fig. S1.**
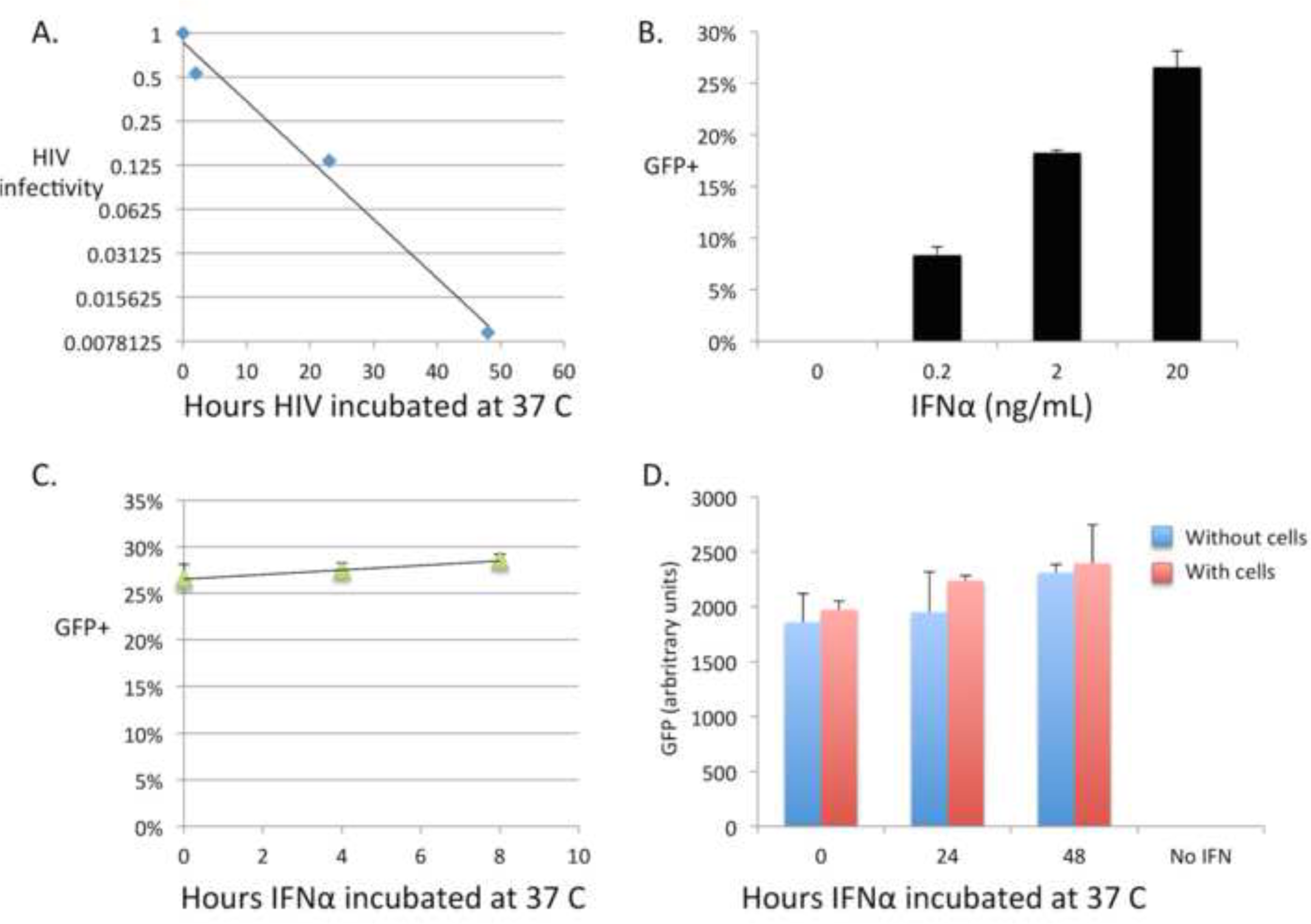

**Fig. S2.**
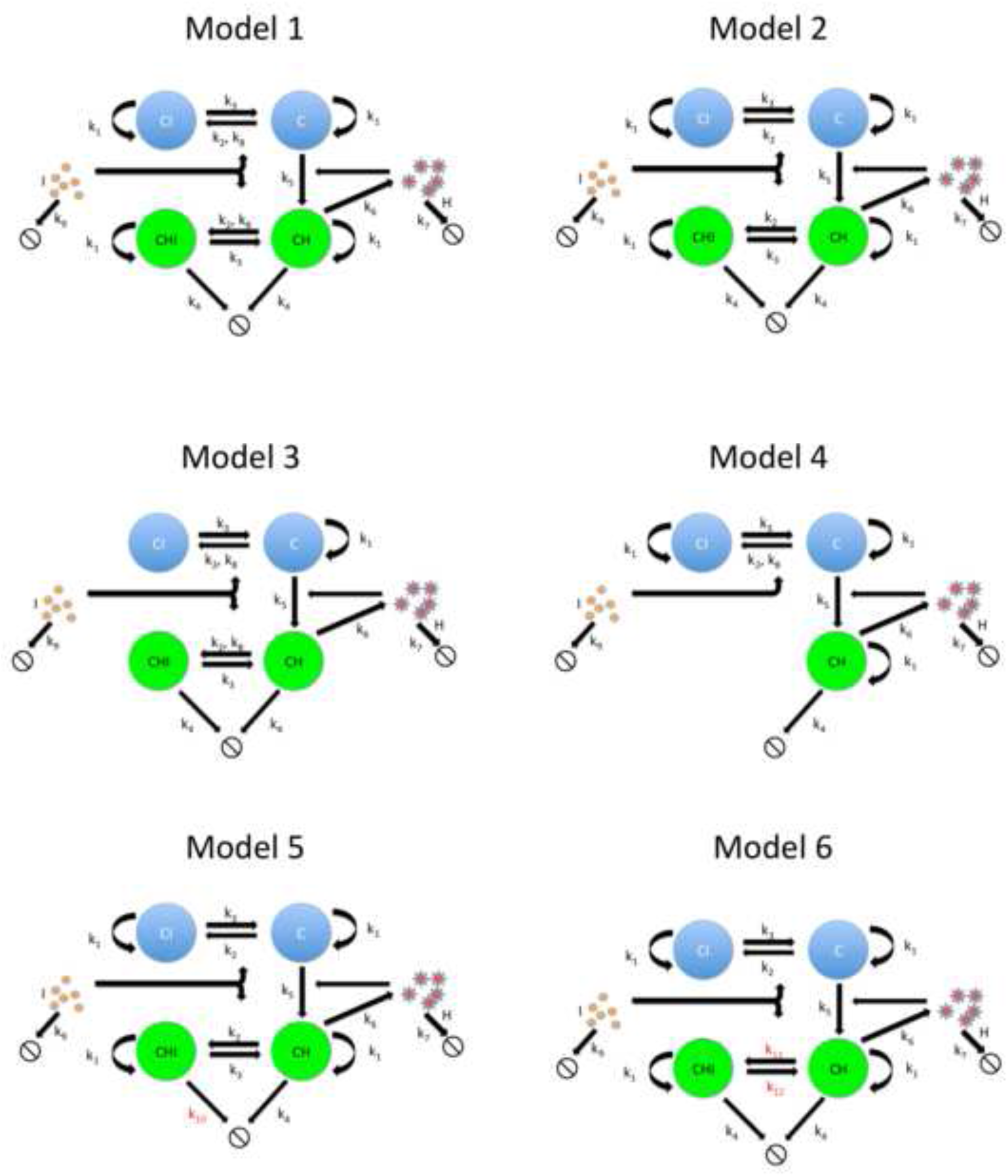

**Fig. S3.**
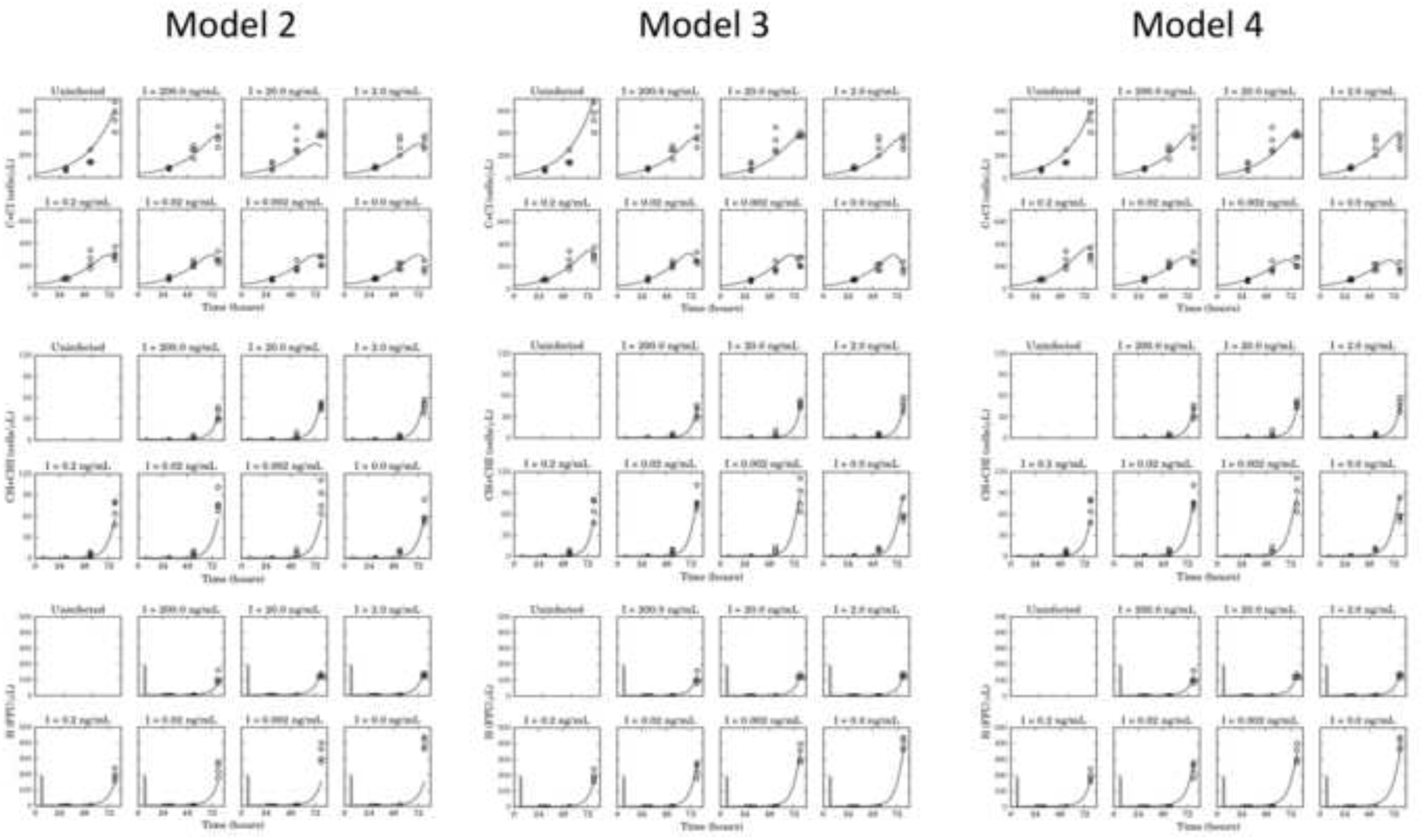

**Fig. S4.**
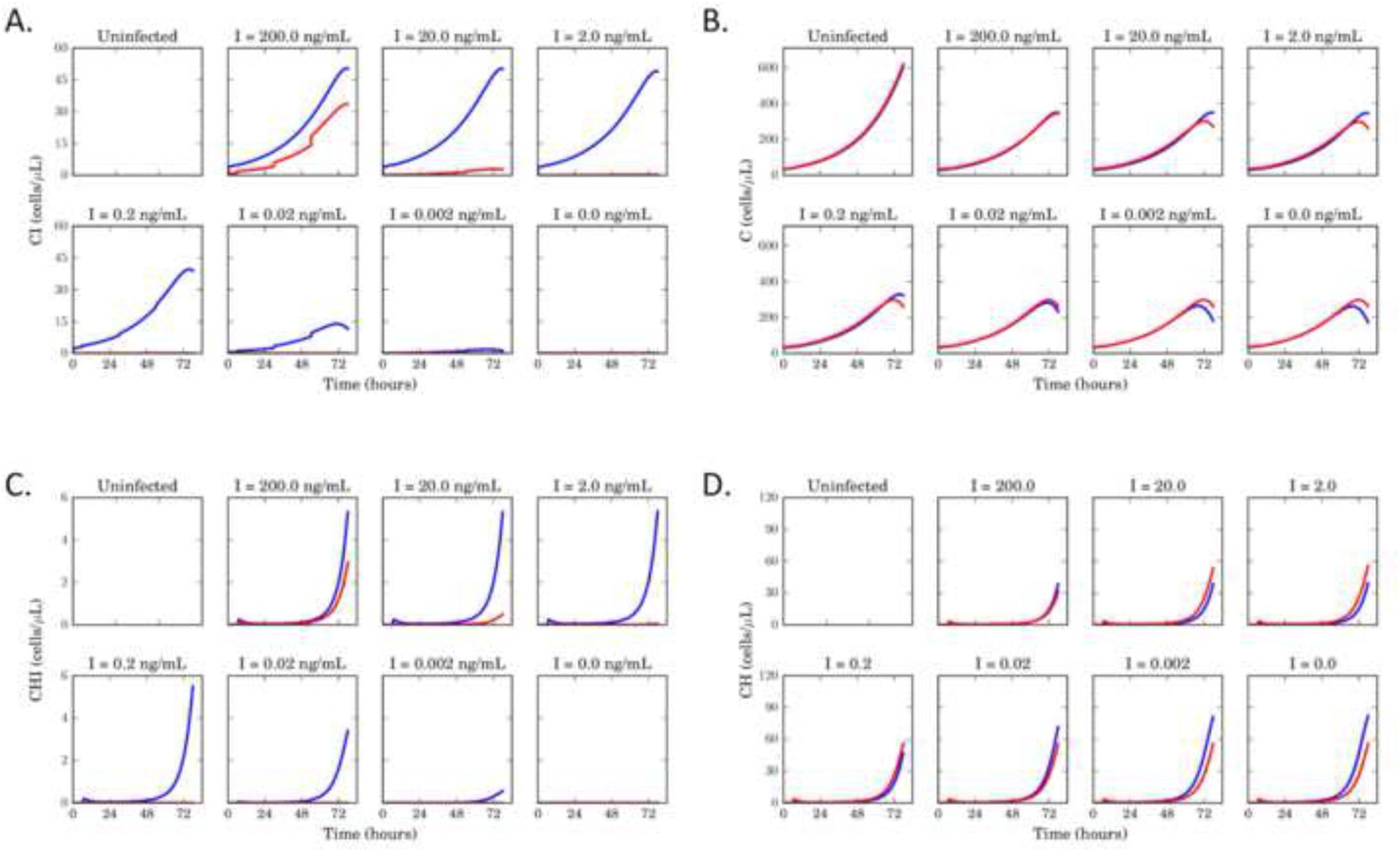

